# Live immunogenic melanoma cells generated by etoposide treatment upregulate B2M and lose tumor-forming potential

**DOI:** 10.64898/2026.10.18.759921

**Authors:** Benjamin W. Lyon, Emma Russo, Cheyenne Colbert, Silas Vega, Tiffany Emmons, Yi-Wen Kong, Annabelle Sanders, Jani Benoit, Hilary Gaudet, Michael B. Yaffe, Ganapathy Sriram

## Abstract

Cytotoxic chemotherapy has been the primary cancer treatment modality for decades, however, chemotherapy drugs do not selectively target cancer cells. These drugs are toxic to all proliferating cells including immune cells that constitute the host-defense system against pathogens and cancer cells. In an effort to develop non-toxic combinations of chemotherapy and immunotherapies for cancer, we previously reported in Sriram et al. 2021 that etoposide-treated B16 melanoma cells expressing ovalbumin, when co-cultured *in vitro* with dendritic cells and T-cells, enhance T-cell activation. The live fraction of melanoma cells after etoposide treatment was critical to promote T-cell activation as dead cells alone or in combination with cell-free supernatants were not effective. In this study, we show that the live fraction of etoposide-treated B16-Ova cells, which gain the ability to stimulate T-cells, are growth-arrested, but remain viable past the co-incubation period with dendritic cells and T-cells post etoposide washout. Furthermore, using a soft-agar colony-formation assay as a proxy for tumor-forming potential, we show that these live etoposide-treated B16-Ova cells have drastically reduced ability to form colonies displaying a near-complete loss of anchorage-independent growth. We also show that etoposide treatment upregulates the production of Beta-2-microglobulin (B2M), a subunit of Class I MHC. Intriguingly, we observed using immunofluorescence that the live etoposide-treated B16-Ova cells show enhanced intracellular perinuclear B2M accumulation but B2M surface staining, while detectable, is not enhanced by etoposide treatment. Furthermore, while the live etoposide-treated B16-Ova cells express Class I MHC protein, there is only a marginal drug treatment-induced increase observed in the expression levels by western blotting. Our results suggest that T-cell activation by live etoposide-treated B16-Ova cells is likely not driven by drug treatment-induced enhancement in cell surface levels of Class I MHC and point to other mechanisms.

## INTRODUCTION

Cytotoxic chemotherapy has been at the core of cancer treatment for decades and continues to be the first line of treatment for many cancer types including colon cancer, lung cancer and pancreatic cancer [1]. While many tumor types show promising initial responses to chemotherapy, relapse of therapy-resistant disease frequently occurs in patients. Notably, most chemotherapy drugs have toxic side-effects, many of which result from the toxicity to actively proliferating non-cancer cells in the body, particularly cells that originate in the bone marrow. Immunotherapy has emerged as a promising new treatment modality wherein targeting host T-cells with the immune checkpoint inhibitors anti-PD1, anti-PD-L1 and/or anti-CTLA-4 evokes remarkable anti-tumor responses in a subset of patients in advanced disease stages, resulting in tumor shrinkage and tumor-free survival for a number of years beyond what was possible with conventional chemotherapy. While this is a breakthrough in cancer treatment, responses are still limited to 10-30% of patients in most tumor types [2, 3]. Combining chemotherapy with immunotherapy has resulted in improved responses in some tumor types [4-7]. However, there is a need to identify chemotherapy drugs and treatment methods that would combine optimally with specific immunotherapies to achieve maximal responses.

We previously published that when specific chemotherapy drugs were used at specific doses to treat B16 melanoma cells (expressing ovalbumin), followed by drug washout and sequential co-incubation with bone marrow-derived dendritic cells and OT-1 CD8+ T-cells, production of IFN-γ by OT-1 CD8+ T-cells was enhanced [8, 9]. Interestingly, the live-attached fraction of tumor cells after chemotherapy treatment was sufficient to promote T-cell IFN-γ production in this assay but the dead cells or the cell-free supernatants alone did not induce T-cell activation. These results indicated that tumor cells were converted into a live immunogenic state after chemotherapy treatment in a drug and dose-dependent manner. Furthermore, we also showed that when B16-Ova tumor cells after treatment with the chemotherapy drug etoposide (followed by drug washout) were directly injected into melanoma tumors in mice, in combination with systemic Immune Checkpoint Inhibition (i.e. anti-PD1 + anti-CTLA-4 antibodies, hereafter referred to as ICI), complete tumor shrinkage and resistance to re-challenge was induced in a subset of mice, which was not observed with ICI alone. The mechanisms by which live etoposide-treated B16-Ova tumor cells stimulate T-cell activation remain to be elucidated. Other studies have also demonstrated the potential of modified cancer cells to re-activate anti-tumor immunity and induce tumor regression in animal models [10-13]. Our study along with these studies raises the possibility of using modified cancer cells as a therapeutic modality. However, for this to be feasible, it would be critical to evaluate if live immunogenic chemotherapy-treated cancer cells still have growth potential and retain the ability to form tumors.

In this study, we investigated the mechanisms by which the live-fraction of etoposide-treated B16-Ova melanoma cells promotes T-cell activation and if this fraction of cells showed growth-arrest and became senescent. We show here that the live fraction of B16-Ova cells after etoposide treatment was growth-arrested but continued to stay viable for at least 120 hours after start of etoposide treatment. Notably, the live etoposide-treated cell fraction shows a near complete loss of soft agar colony formation ability indicating loss of tumor-forming potential. We found that the live-attached fraction of etoposide-treated B16-Ova cells that are sufficient to stimulate T-cell activation do not display any substantial beta-galactosidase activity 48 hours after start of treatment. We also show that etoposide treatment upregulates the expression of B2M protein, but not mRNA, in B16-Ova cells. B2M levels are enhanced in the perinuclear region of etoposide-treated B16-Ova cells, however, while punctate staining characteristic of surface expression is detectable, there is no treatment-induced enhancement suggesting that B2M expression is not enhanced on the cell surface. Total MHC I protein levels are detectable in the live etoposide-treated B16-Ova cell fraction but only marginally increased compared to the DMSO control as observed by western blotting compared to the strong upregulation observed by treatment of cells with IFN-γ. Our results indicate that enhanced B2M expression induced by etoposide does not manifest as substantially enhanced cell surface MHC I levels pointing at alternate mechanisms by which live etoposide-treated B16-Ova cells promote T-cell activation.

## METHODS

### Cell culture and reagents

B16-Ova murine melanoma cells were maintained in Dulbecco’s Modified Eagle Medium (DMEM) supplemented with 10% fetal bovine serum (FBS) and 1% GlutaMAX. The following antibodies were purchased from Cell Signaling Technology Inc.: Anti-beta-2-microglobulin (B2M) (cat #64046), Anti Beta-Actin (Cat #3700), anti-tapasin (Cat # 66382T), anti-H2D1 (Class I MHC, cat # 35923T). Beta-galactosidase staining kit was also purchased from Cell Signaling Technology (Cat # 9860S). Anti-p21 (Cat # MA569765), ProLong Gold Antifade Mountant with DNA Stain DAPI (Cat #P36941) and TriZol (Cat # 15596026) was purchased from Invitrogen. Alexa Fluor 488 AffiniPure Goat Anti-Rabbit IgG (Cat #111-545-003) and normal goat serum (Cat # 005000121) was purchased from Jackson Immunoresearch Labs. Cell Titer glo kit was purchased from Promega (Cat # G7570). RevertAid cDNA synthesis kit was purchased from Thermo scientific (Cat # K1621). Recombinant mouse IFN-γ was purchased from Biolegend (Cat # 575304). Etoposide was purchased from Sigma (Cat # E1383 – 100MG).

### Viability assays

To assess cell viability, B16-Ova cells were treated for 24 hours with the drug compounds followed by drug and dead cell washout. The live-attached fraction of cells was harvested and 5,000 cells were plated per well in a white opaque 96-well plate with three replicates per condition per time point. At each measured time point post-treatment (excluding 24 hours), the media was removed and replaced with 75μL fresh media at room temperature. 25μL CellTiter-Glo reagent was added to wells and incubated at room temperature for 20-30 minutes. Cells measured at 24 hours post-treatment were plated in 75μL medium and 25μL CellTiter-Glo reagent was added immediately followed by incubation at room temperature for 20-30 minutes. Luminescence was read on a BioTek Synergy H1 microplate reader.

### Senescence assay

This assay was performed according to the manufacturer’s instructions in the beta-galactosidase staining kit purchased from Cell Signaling Technology (Cat # 9860S).

### Soft agar colony formation assay

To evaluate anchorage-independent growth of the live-attached fraction of chemotherapy-treated melanoma cells, a soft agar colony formation assay was performed. A solid bottom layer was added to culture dishes using a 0.8% agarose solution to prevent cell attachment. Once the base layer solidified, the live-attached fraction of B16-OVA cells after chemotherapy treatment was harvested and suspended in a semi-solid matrix consisting of 0.4% agarose and culture media. 5,000 cells were suspended per dish, with triplicates for each condition.

### Fluorescence microscopy

Twenty-millimeter circular glass coverslips were placed in 10 cm dishes and sterilized via microwave for 30 seconds. B16-OVA cells (1,000,000 per dish) were seeded and allowed to adhere for 24 hours. Cells were then treated with DMSO (control), etoposide (50 μM), or IFN-γ (40 ng/uL). After 24 hours, the drugs and dead cells were washed out and the live-attached fraction of cells was washed three times with 1x PBS and fixed with 4% paraformaldehyde (PFA) for 15 minutes at room temperature. After three subsequent PBS washes, cells were blocked and permeabilized for 45 minutes in 1x PBS containing 2.5% normal goat serum and 0.1% Triton X-100. Primary antibody incubation was performed at 4 °C for 18 hours using the anti-B2M antibody diluted 1:3000 in blocking buffer. Secondary antibody incubation was performed using goat anti-rabbit Alexa Fluor 488 at a 1:750 dilution for 1.5 hours at room temperature, also in blocking buffer. Coverslips were mounted using ProLong Gold Antifade mountant with DAPI. Images were acquired at 40x magnification. To facilitate quantitative comparison, all images within a series were captured using identical exposure settings.

### Quantification of fluorescence imaging

Image processing was performed using ImageJ (NIH). All in-focus cells within a field were selected for quantification. The B2M and DAPI channels were merged to define cellular boundaries, and individual cell outlines were drawn using the polygon selection tool. To isolate the cytoplasmic versus surface signal, the perinuclear B2M region was manually defined as a Region of Interest (ROI). In cells lacking a distinct perinuclear accumulation, an ROI was placed in the expected anatomical location. The perinuclear ROI was subtracted from the total cellular ROI to segment the peripheral signal. The area-normalized integrated density was recorded for each cell from each coverslip and plotted.

### RT-qPCR

B16-Ova cells (1,000,000) were seeded in 10 cm dishes and treated with etoposide or IFN-γ. After 24 hours, the live-attached fraction of cells was harvested via trypsinization and pelleted at 200 x g for 5 minutes. Total RNA was extracted using the TRIzol Reagent according to the manufacturer’s instructions. cDNA synthesis was performed using the Thermo Scientific RevertAid First Strand cDNA Synthesis Kit. Fisherbrand SYBR green qPCR master mix was used. The following primer sequences were used:

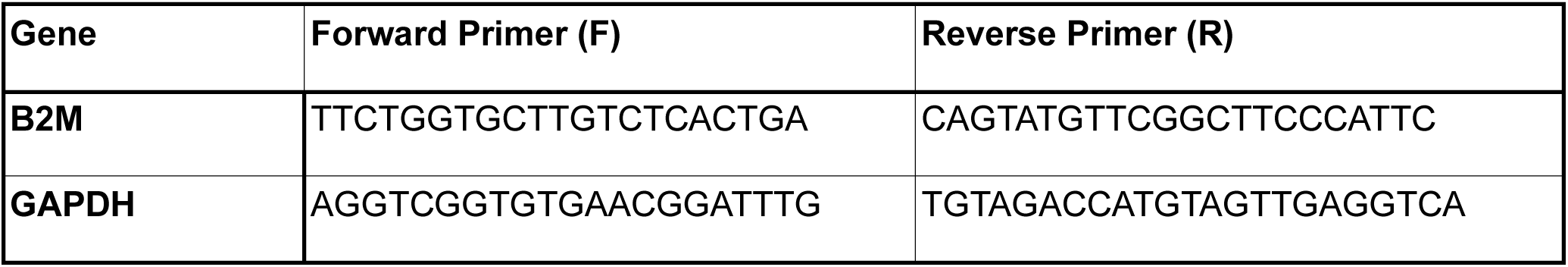

### Statistical analysis

All statistical analyses were performed using Graphpad PRISM. Comparisons between three or more treatment conditions were performed using ANOVA followed by Dunnett’s or Sidak’s multiple comparisons tests post-hoc as indicated in the figure legend. Comparisons between two treatment conditions were performed using a two-tailed t-test.

## RESULTS

As shown in Sriram et al. 2021, etoposide-treated B16-Ova murine melanoma cells promote T-cell activation in a co-culture assay wherein B16-Ova cells are treated with etoposide followed by drug washout, and co-incubated with dendritic cells followed by addition of OT-1 CD8+ T-cells [8]. Furthermore, etoposide-treated B16-Ova cells, when injected directly into melanoma tumors in mice induce complete tumor regressions and immunological memory in a subset of mice in combination with systemic anti-PD-1 and anti-CTLA-4 antibodies. These results indicate that etoposide-induced cell injury in B16-Ova cells is immunogenic, serving as a stimulus for the cancer-immunity cycle.

### Live attached B16-Ova cells but not dead cells generated after etoposide treatment promote T-cell activation

To test and demonstrate the relative contributions of the live-attached B16-Ova cell fraction and the dead floating cell fraction, as shown previously in Sriram et al. 2021, B16-Ova cells were treated with etoposide at 50 μM for 24 hours followed by drug washout and harvesting of the total treated cell fraction (live+dead), the live-attached cell fraction only and the dead (floating) cell fraction. Each of the above fractions were co-incubated separately with murine bone marrow-derived dendritic cells for 24 hours followed by addition of OT-1 CD8+ T-cells. IFN-γ production by T-cells was then assessed by intracellular cytokine staining and flow cytometry (Fig 1A). As shown in Figure 1B, the total (live+dead) fraction of B16-Ova cells after treatment with etoposide at 50 μM stimulates IFN-γ production in OT-1 T-cells. Notably, as shown previously in Sriram et al. 2021, the live-attached fraction of etoposide-treated B16-Ova cells but not the dead floating cell fraction stimulates T-cell IFN-γ production indicating that the live attached fraction of B16-Ova cells after etoposide treatment is sufficient to stimulate T-cell activation.

**Figure 1:**
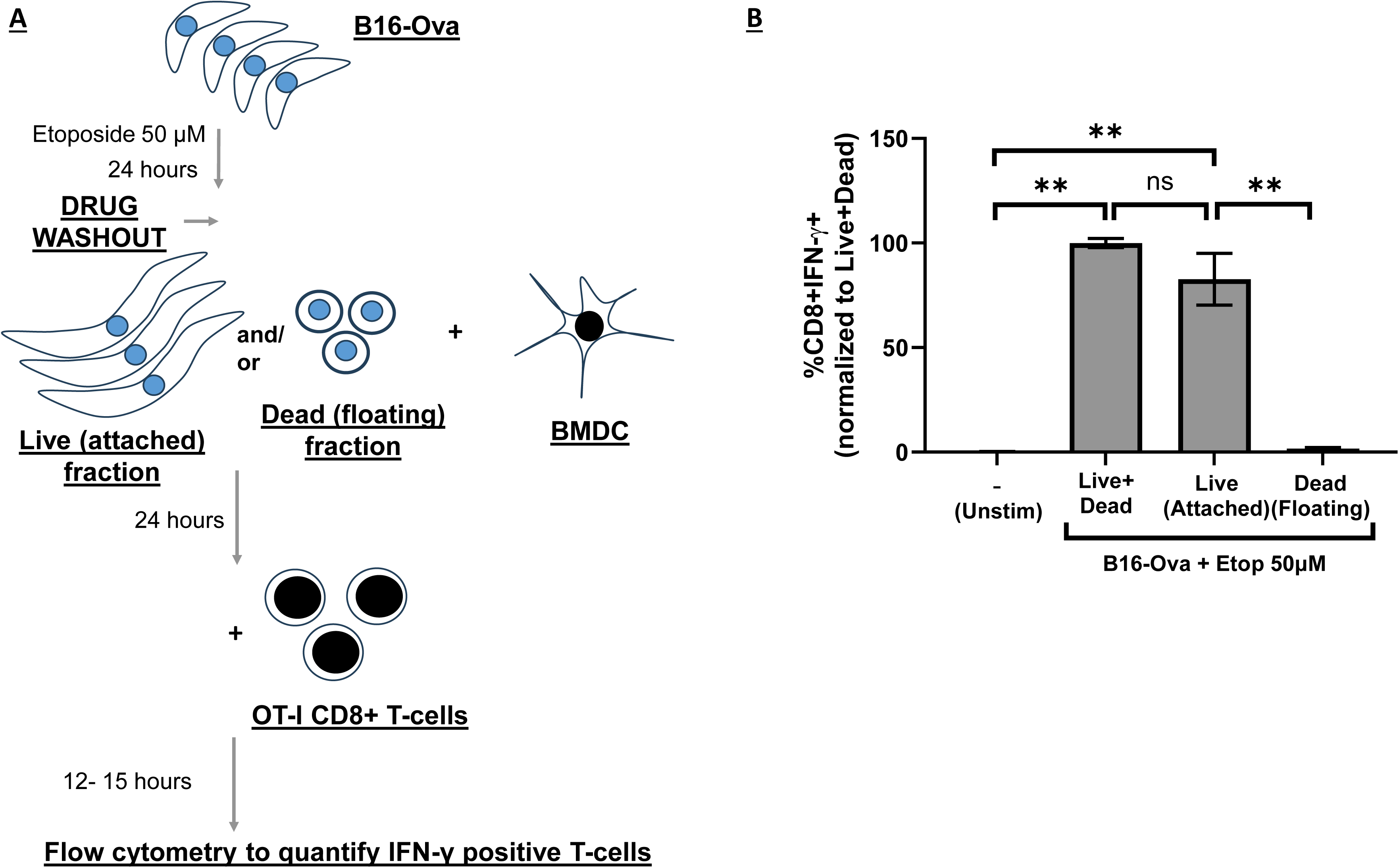
Live-attached B16-Ova cells generated by etoposide treatment but not dead (floating) cells promote T-cell activation. **A.** Schematic showing the co-culture assay wherein B16-Ova were treated with etoposide or DMSO for 24 hours, drug washout was done and either the total treated B16-Ova cell fraction (Live+dead) or the live attached B16-Ova cell fraction after etoposide treatment or the dead (floating) cell fraction were separately co-cultured with bone marrow-derived dendritic cells for 24 hours. This was followed by the addition of OT-1 CD8+ T-cells and 12-15 hours later, IFN-γ positive T-cells were determined by intracellular cytokine staining and flow cytometry. “- (Unstim)“ refers to BMDC and OT-1 CD8+ T-cells co-cultured in the absence of B16-Ova cells. **B.** Quantification of IFN-γ+ OT-1 CD8+ T-cells is shown as mean +/- SEM. “**” indicates p<0.05 using ANOVA followed by Sidak’s multiple comparisons test. Results are representative of two independent experiments. All values are expressed as a percentage of IFN-γ+ T-cells observed in the (Live+dead) group.

### The live-attached fraction of B16-Ova cells generated after etoposide treatment are alive for up to 5 days after start of treatment but appear to be growth-arrested

To examine the morphology and to track the live-attached fraction of B16-Ova tumor cells after chemotherapy treatment beyond 24 hours of treatment, B16-Ova cells were treated for 24 hours with etoposide at 50 μM, drugs and dead cells were washed out and the live-attached fraction of cells was imaged by phase contrast microscopy every 24 hours (as outlined in Fig 2A). As shown in the representative images in Figure 2B, the live-attached fraction of cells after etoposide-treatment remains attached and cells are at nearly the same confluence (visually) over 120 hours. The live-attached etoposide-treated fraction of cells frequently shows elongated morphology typical of stressed cells compared to the control (DMSO-treated cells).

**Figure 2:**
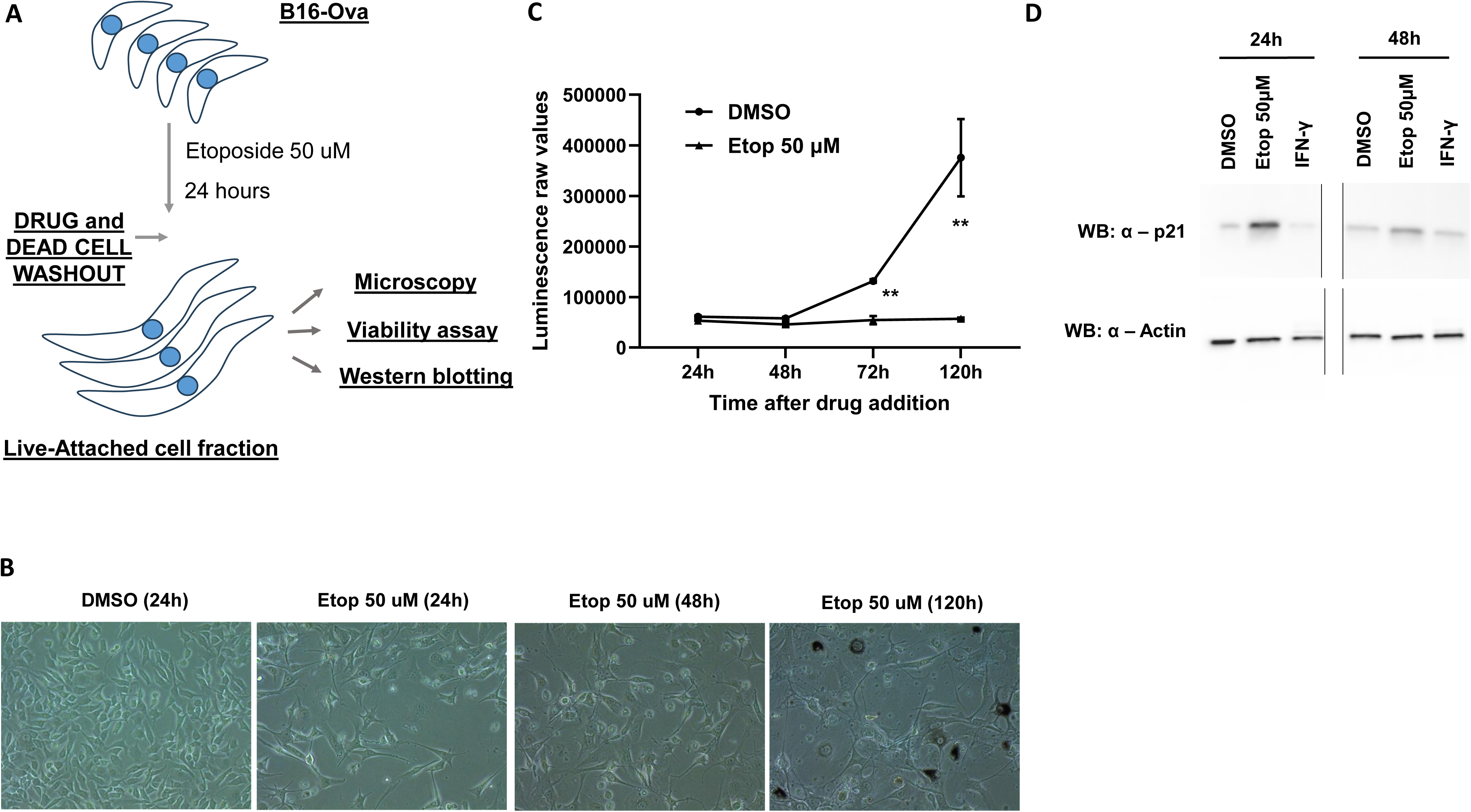
Live etoposide-treated B16-Ova cells upregulate p21 and are growth arrested but remain viable for at least 96 hours post drug washout. **A.** Schematic showing how the live-attached fraction of B16-Ova was obtained for analysis after 24 hours of treatment with etoposide followed by drug washout. **B.** Phase contrast images of the live-attached fraction of B16-Ova cells obtained as outlined in A. **C.** Quantification of luminescence values obtained from a Cell Titer Glo assay on the live attached fraction of B16-Ova cells obtained as outlined in A at the indicated time points after start of treatment. At each time point, luminescence is depicted as mean +/- SEM. Results are representative of two independent experiments. ‘**’ indicates p<0.05 using a two-way ANOVA followed by Sidak’s multiple comparisons test. **D.** Western blots for p21 and beta-actin are shown on the live attached fraction of B16-Ova cells harvested immediately after treatment for 24 hours and drug washout or harvested after an additional 24 hours after drug washout (total 48 h after start of treatment). Results are representative of two independent experiments.

To quantify viability of the live-attached fraction of B16-Ova cells after treatment with chemotherapy, cells were treated with etoposide at 50 μM for 24 hours followed by drug washout and re-plating of the attached fraction into white opaque 96-well plates for the cell-titer glo assay. Luminescence was measured at 24 hours, 48 hours, 72 hours and 120 hours after start of treatment. As shown in Figure 2C, cells treated with DMSO show a steady increase in luminescence at each measured time point, indicative of cell proliferation. Conversely, cells treated with 50 μM etoposide show significant growth inhibition as indicated by the luminescence levels remaining near baseline throughout the 120-hour period. These results indicate that etoposide treatment at 50 μM induces a state of growth-arrest in the attached fraction of B16-Ova cells, but these cells continue to stay alive for at least 120 hours after the start of treatment.

### Etoposide-treated B16-Ova cells show enhanced p21 levels

To investigate this state of cell cycle arrest observed in the live-attached etoposide-treated B16-Ova cell fraction further, the expression of the CDK inhibitor p21 was evaluated. B16-Ova cells were treated with DMSO or etoposide (50 μM) for 24 hours. After 24 hours, the drug was washed out and the live-attached fraction of cells was either immediately harvested or incubated in drug-free media for another 24 hours and then harvested. Cell lysates were analyzed by western blotting as shown in Figure 2D. p21 was upregulated in the attached fraction of B16-Ova cells at 24 hours after start of treatment and the levels were higher than the DMSO control even at 48 hours. These results are consistent with data shown in Fig 2C and indicate that the live-attached fraction of B16-Ova cells generated after treatment with etoposide at 50 μM are in a state of cell-cycle arrest.

### B16-Ova cells treated with etoposide at 50 μM are not beta-galactosidase-positive at 48 hours

To investigate if this state of cell cycle arrest results in senescence, B16-Ova cells were treated with DMSO or etoposide at 50 μM for 24 hours after which drugs were washed out and fresh drug-free media was added. Cells were fixed at 48 hours and 144h after the start of drug treatment followed by staining for beta-galactosidase activity. As shown in the representative images in Fig 3A, at 48 h post start of treatment, no beta-galactosidase-positive cells were observed in the controls or the live-attached etoposide-treated fraction of B16-Ova cells as indicated by the absence of any blue staining. At 144h post start of treatment, no beta-galactosidase-positive cells are observed in the DMSO-treated controls but a few are observed in the live etoposide-treated fraction (black arrows in Fig 3A) indicating a small minority of B16-Ova cells are senescent at this time point (see quantification in the right panel of Fig 3A). As shown in Fig 1, immunogenicity of etoposide-treated B16-Ova cells is assessed by initial co-incubation with DCs starting at 24 hours after etoposide treatment of B16-Ova cells for a further 24 hours. At this time point i.e. 48 hours after start of etoposide exposure of B16-Ova cells is when OT-1 CD8+ T-cells are added to the co-culture. As no beta-galactosidase-positive B16-Ova cells are observed at this point, these results suggest that the gain of immunogenicity by the live etoposide-treated B16-Ova cells precedes the onset of classical senescence markers.

**Figure 3:**
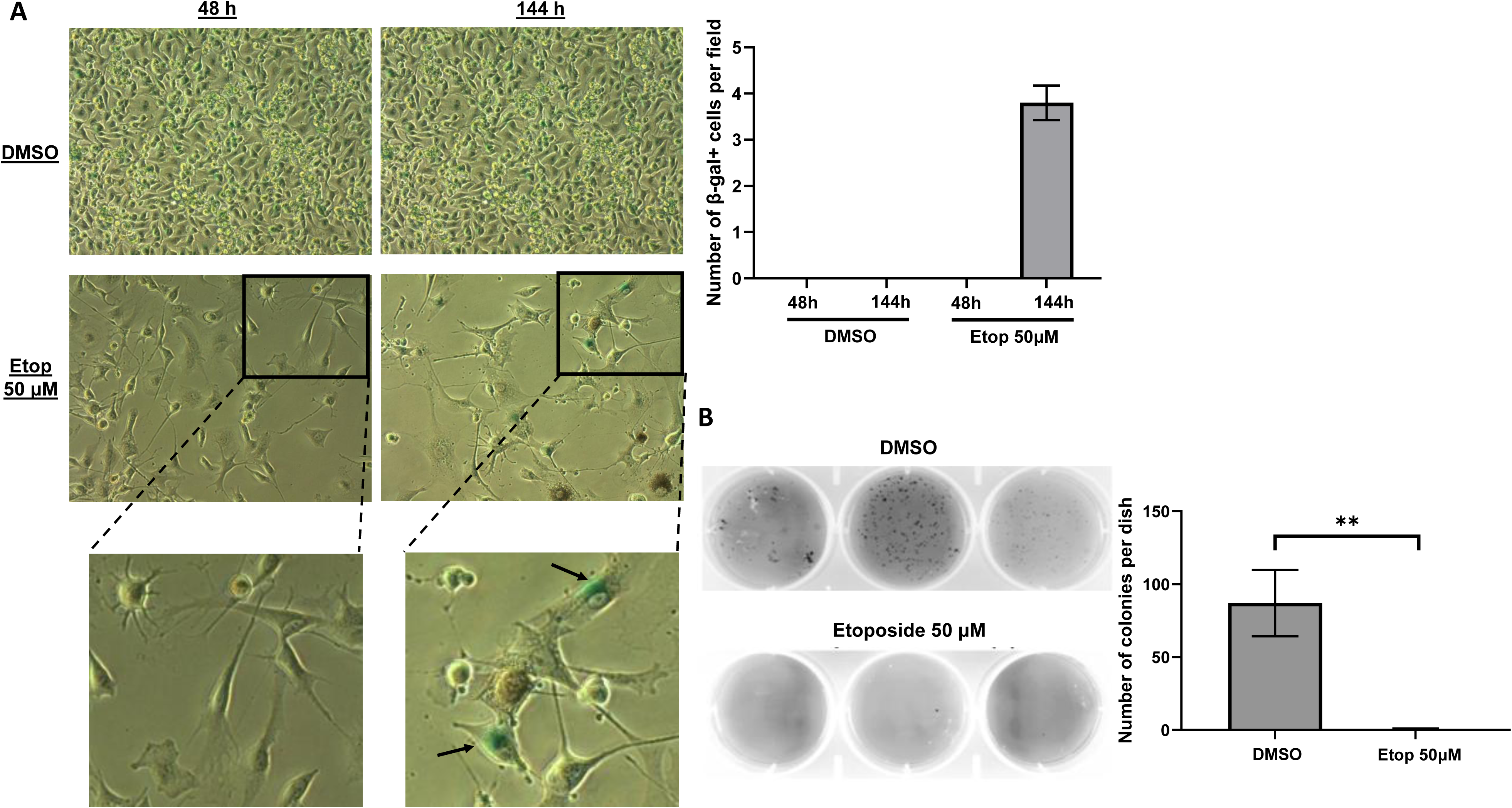
Live etoposide-treated B16-Ova cells are not beta-galactosidase-positive at 48 hours after start of treatment and show drastically reduced ability to form colonies on soft agar. **A.** Beta-galactosidase assay performed on the live-attached fraction of B16-Ova cells after treatment with etoposide at 50 μM for 24 h followed by drug and dead cell washout and addition of fresh media for a further 24h (labeled 48 h) or 120 h (labeled 144 h). Representative phase contrast images are shown. Results are representative of two independent experiments. Quantification of beta-galactosidase-positive (blue) cells is shown on the right. Arrows in the enlarged sub-sections point to beta-galactosidase-positive cells. **B.** Soft-agar colony formation assay was performed on the live-attached fraction of B16-Ova cells obtained after treatment with etoposide at 50 μM for 24h. Colony formation was assessed 21 days after seeding in soft agar. Images are shown after crystal violet staining. Results are representative of three independent experiments. Quantification is shown on the right as mean +/- SEM. ‘**’ indicates p<.05 using a two-tailed t-test.

### Evaluating tumor forming ability of etoposide-treated melanoma cells

To evaluate the tumor-forming potential of the live-attached fraction of B16-Ova cells generated by etoposide treatment, a soft agar colony formation assay was utilized to assess whether etoposide-treated cells maintain anchorage-independent growth. B16-Ova cells were treated with DMSO or 50μM etoposide for 24 hours. The drugs were removed and the attached fraction of cells was harvested, washed and re-suspended in a layer of 0.4% agarose-containing media and cast in a tissue culture dish on top of a 0.8% agarose-containing solid layer. Plates were incubated for 21 days before colony formation was assessed by staining with crystal violet. As shown in Figure 3B, DMSO-treated cells show significant anchorage-independent proliferation and high-density colonies demonstrating the tumor-forming potential of the B16-Ova cell line. Contrastingly, 50μM etoposide treatment resulted in near-complete elimination of colony formation. These results indicate that the live-attached fraction of B16-Ova cells generated by etoposide treatment show a near-complete loss of anchorage-independent growth indicating that this cell fraction is likely no longer tumorigenic.

### Treatment with etoposide enhances expression of beta-2-microglobulin protein

To investigate if etoposide treatment enhanced class I MHC levels in the live-attached fraction of B16-Ova cells, expression of Beta-2-microglobulin (B2M), a critical component of class I MHC, was assessed. B16-Ova cells were treated with DMSO, etoposide or IFN-γ for 24 hours, following which the drug and dead cells were washed out and the live-attached cell fractions were harvested. Analysis of B2M mRNA levels was performed by qPCR and protein levels by SDS-PAGE and western blotting of the lysates of the live-attached cell fractions. As shown in Figure 4A, treatment with etoposide did not significantly upregulate B2M mRNA levels in the live-attached B16-Ova cell fraction compared to the DMSO control. On the other hand, treatment with IFN-γ strongly upregulated B2M mRNA levels. Interestingly, as shown in Figure 4B, while IFN-γ treatment strongly enhanced B2M protein levels compared to the DMSO control, enhanced protein levels of B2M were also observed in cells treated with 50 μM etoposide but not with 10 μM etoposide compared to the DMSO-treated control. These results prompted us to investigate the surface levels of B2M as a proxy for Class I MHC expression on the live-attached fraction of etoposide-treated B16-Ova cells.

**Figure 4:**
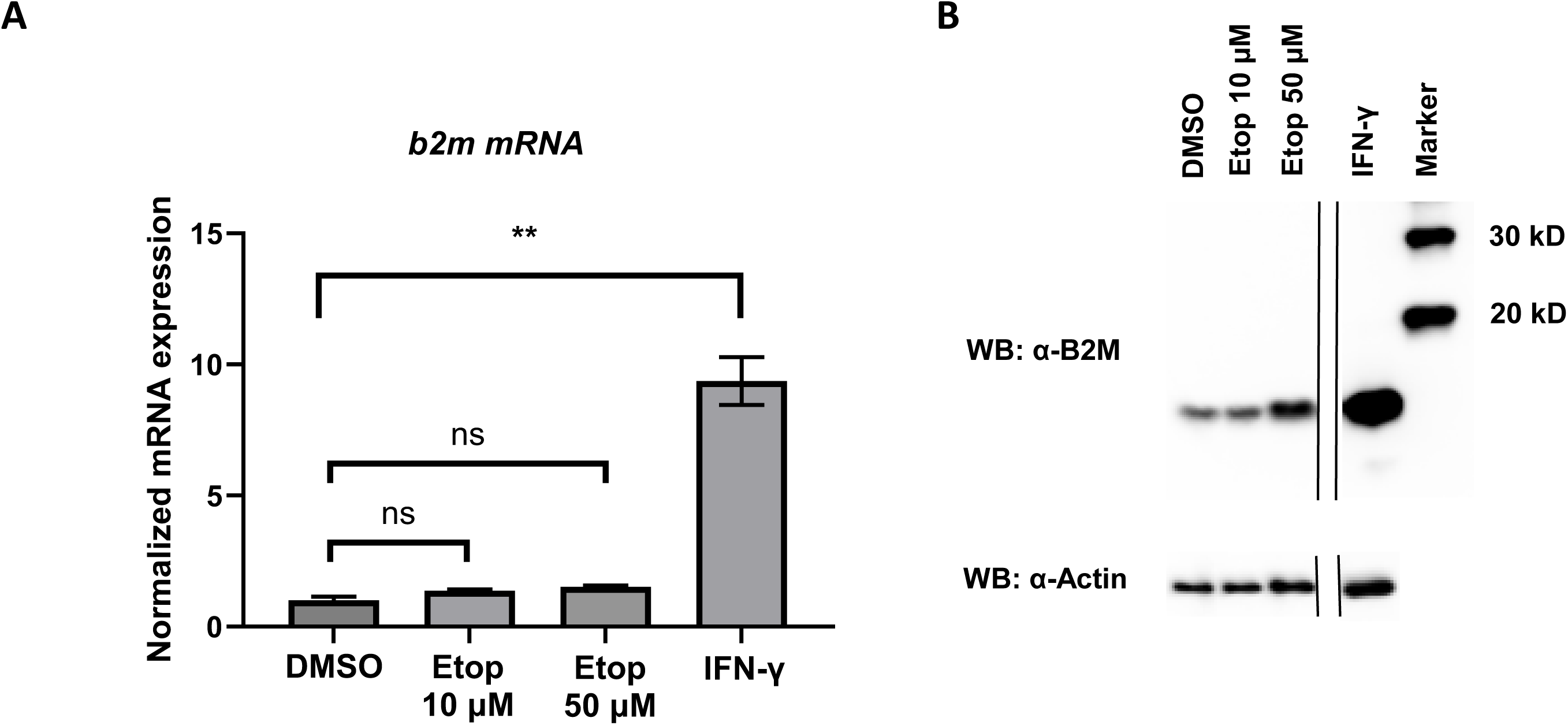
Etoposide treatment upregulates the expression of Beta-2-microglobulin (B2M) in B16-Ova cells. **A.** B16-Ova cells were treated with DMSO, etoposide or IFN-γ for 24 hours followed by drug washout. The live-attached fraction was harvested, total RNA was extracted, cDNA was generated and qPCR was performed to assess the mRNA levels of B2M using GAPDH as a control. Quantification is shown as mean +/- SEM. ‘**’ indicates p<.05 using ANOVA followed by Dunnett’s multiple comparisons test. **B.** B16-Ova cells were treated with DMSO, etoposide or IFN-γ for 24 hours followed by drug washout. The live-attached fraction was harvested, cell lysates were generated and analyzed by SDS-PAGE and western blotting with anti-B2M and anti-beta-actin antibodies. Results are representative of two independent experiments.

### Etoposide treatment enhances perinuclear staining but not punctate staining in B16-Ova cells

To test if surface Class I MHC levels were enhanced in etoposide-treated B16-Ova cells, immunofluorescence was performed using an anti-B2M antibody (as a proxy for surface class I MHC levels). B16-Ova cells were plated on coverslips and then treated with etoposide at 50 μM for 24 hours. Treatment with IFN-γ was used as a positive control. After treatment, the drugs and dead cells were washed out and the live-attached fraction of cells was fixed, and staining was performed with an anti-B2M primary antibody and goat anti-rabbit Alexa Fluor 488 conjugated secondary antibody.

As shown in Figure 5A (Compare DMSO and IFN-γ panel), IFN-γ-treated B16-Ova cells displayed enhanced perinuclear staining as well as enhanced punctate staining of B2M compared to the DMSO control, suggesting that there was enhanced intracellular B2M production likely in the endoplasmic reticulum (ER) as well as enhanced cell surface accumulation of B2M after IFN-γ treatment (quantified in Figs 5B and 5C). On the other hand, the live etoposide-treated B16-Ova cells also displayed enhanced intracellular perinuclear staining and while punctate staining characteristic of surface expression was detectable, it was not enhanced after etoposide treatment (Figures 5D and 5E), which suggested that intracellular B2M production was enhanced, but B2M expression on the surface of cells was altered only minimally or not at all by etoposide treatment.

**Figure 5:**
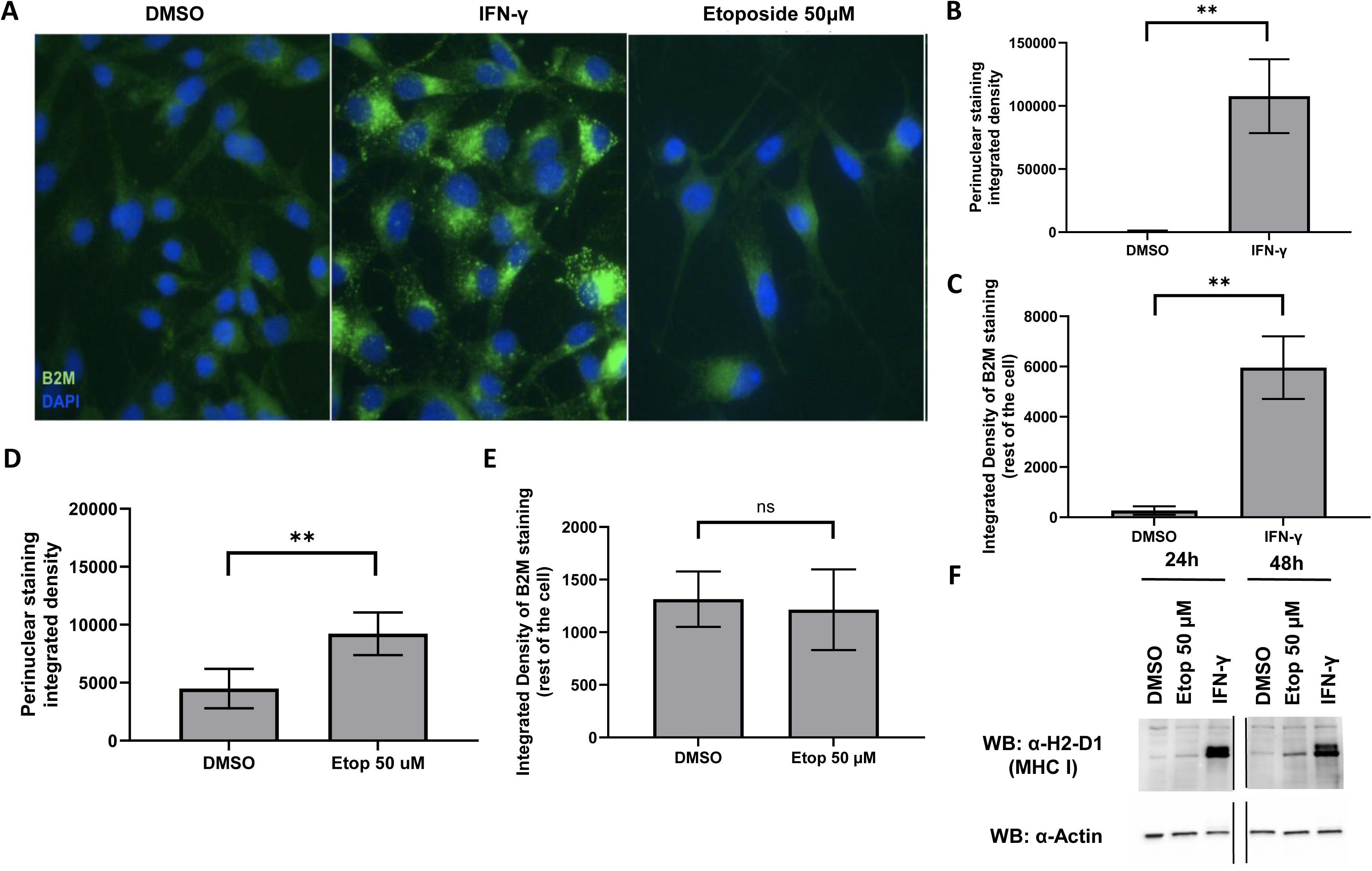
Live etoposide-treated B16-Ova cells show enhanced perinuclear staining of B2M but do not show enhanced B2M punctate staining. **A.** Representative immunofluorescence images of B16-Ova cells treated with DMSO, IFN-γ (40ng/uL) or etoposide (50uM). **B.** Quantification of integrated density of perinuclear B2M staining on the live-attached fraction of B16-Ova cells obtained after treatment with DMSO or IFN-γ for 24 hours. Quantification is shown as mean +/- SEM. Number of cells analyzed for DMSO = 39, IFN-γ = 29. ‘**’ indicates p< 0.05 using a two-tailed t-test. **C.** Quantification of integrated density of punctate B2M staining on the live-attached fraction of B16-Ova cells obtained after treatment with DMSO or IFN-γ for 24 hours. Quantification is shown as mean +/- SEM. Number of cells analyzed for DMSO = 39, IFN-γ = 29. ‘**’ indicates p< 0.05 using a two-tailed t-test. **D.** Quantification of integrated density of perinuclear B2M staining on the live-attached fraction of B16-Ova cells obtained after DMSO or etoposide (50 μM) treatment for 24 hours of is shown as mean +/- SEM. Number of cells analyzed for DMSO = 132, IFN-γ = 142. ‘**’ indicates p< 0.05 using a two-tailed t-test. **E.** Quantification of integrated density of punctate B2M staining on the live-attached fraction of B16-Ova cells obtained after DMSO or etoposide (50 μM) treatment for 24 hours is shown as mean +/- SEM. Number of cells analyzed for DMSO = 132, IFN-γ = 142. ‘**’ indicates p< 0.05 using a two-tailed t-test. **G.** B16-Ova cells were treated with DMSO, etoposide or IFN-γ for 24 hours followed by drug and dead cell washout. The live-attached cell fraction was harvested, cell lysates were generated and analyzed by SDS-PAGE and western blotting with anti-MHC I and anti-beta-actin antibodies. Results are representative of two independent experiments.

### Live etoposide-treated B16-Ova cells show only a marginal increase in total MHC I

We next assessed total MHC I protein levels in cells after etoposide treatment by SDS-PAGE and western blotting. B16-Ova cells were treated with DMSO, IFN-γ or etoposide (50 μM) for 24 hours followed by washout of the drugs and the floating cell fraction. The live-attached fraction of cells was either harvested at this time point or incubated in drug-free media for a further 24 hours and then harvested at the 48-hour time-point. Lysates of the live-attached cell fraction at each time point were analyzed by western blotting for total MHC I and beta-actin. As shown in Figure 5F, total MHC I is strongly upregulated in B16-Ova cells after IFN-γ treatment at both 24 hours and 48 hours. In stark contrast, while MHC I is detectable in the DMSO control, no clear treatment-induced enhancement in total MHC I is observed at 24h in the live-attached etoposide-treated B16-Ova cell fraction, and a weak enhancement compared to IFN-γ-treated cells is observed at the 48h time point. In summary, our results indicate that while B16-Ova cells express detectable levels of Class I MHC at baseline, drug treatment-induced increase in total B2M protein expression in the live-attached fraction of etoposide-treated B16-Ova cells does not manifest in a strong enhancement in surface MHC I levels pointing to other mechanisms that may contribute to T-cell activation.

## DISCUSSION

We have previously shown (in Sriram et al 2021) that B16-Ova cells treated with specific doses of doxorubicin, etoposide or mitoxantrone, when co-cultured with bone marrow-derived dendritic cells and OT-1 CD8+ T-cells, promote T-cell activation. The live-attached fraction of chemotherapy-treated B16-Ova cells was sufficient to promote T-cell activation but the dead cell fraction or the cell-free supernatants alone did not promote T-cell activation. While the ability of etoposide-treated B16-Ova cells to promote T-cell activation was dependent on DNA-dependent protein kinase and NF-KB signaling, the contribution of surface antigen presentation and senescence in the live tumor cell fraction after etoposide treatment is not clear. Furthermore, the ability of live etoposide-treated B16-Ova cells to grow and their tumor-forming potential has not been evaluated.

In this study, we show that just as previously reported in Sriram et al 2021, the live-attached fraction of etoposide-treated B16-Ova cells but not the dead floating cell fraction promotes T-cell activation. The live fraction of etoposide-treated B16-Ova cells shows a near complete loss of tumor-forming potential and is growth-arrested. However, this live immunogenic fraction is not beta-galactosidase-positive at 48 hours after start of treatment indicating that the gain of T-cell-activating potential of etoposide-treated B16-Ova cells precedes the onset of this classical senescence marker. The live etoposide-treated B16-Ova cell fraction shows upregulated total B2M protein expression but only a marginal increase in total MHC I levels. Consistently, while punctate B2M staining attributable to surface B2M expression is detectable, there was little to no enhancement in surface B2M staining in the live-attached fraction of etoposide-treated B16-Ova cells even though intracellular perinuclear B2M staining was enhanced. These results indicate that while Class I MHC is detectable at baseline and shows a marginal increase in the live etoposide-treated cells at 48 h, the ability of live etoposide-treated B16-Ova cells to promote T-cell activation in large part is not attributable to treatment-induced increase in MHC I on the B16-Ova cell surface.

Other chemotherapeutic agents including doxorubicin, paclitaxel and bleomycin have been shown to upregulate B2M expression in cancer cells at the mRNA and/or protein levels [14-18]. Each of the above three compounds induced a concomitant increase in MHC I levels in the treated cancer cell population. The effects observed in our study of etoposide on B16-Ova melanoma cells appear to differ from those documented above. B2M is strongly upregulated in etoposide-treated B16-Ova cells as assessed by both western blotting and immunofluorescence but no enhancement in surface B2M is observed and only a marginal increase in total MHC I levels is observed. Given this observation, it is highly likely that B2M is being secreted but this remains to be confirmed experimentally. Whether secreted B2M impacts T-cell activation in this context also remains to be assessed. Additionally, in the absence of substantial etoposide-induced MHC I upregulation on the B16-Ova cell surface, multiple other mechanisms could contribute to enhanced antigen presentation in the live fraction of etoposide-treated B16-Ova cells including enhanced levels of TAP1, TAP2 or tapasin proteins which would result in enhanced antigen loading onto class I MHC. These possibilities need to be evaluated in future work.

The mechanism by which dendritic cells contribute to the T-cell-activating response initiated by the live-fraction etoposide-treated B16-Ova cells is intriguing. Our result implies that DCs potentially receive antigens from live etoposide-treated tumor cells. One possible mechanism for antigen acquisition is through a fraction of live-injured tumor cells that become apoptotic following etoposide treatment. After cell death, DCs engulf apoptotic fragments and acquire tumor antigens which they could subsequently present to T-cells. Another possibility is that the live fraction of B16-Ova cells after etoposide treatment are secreting either soluble antigens or antigens in small membranous vesicles that DCs then acquire [19]. All these possibilities need to be evaluated in future work.

Past work has reported that cancer cells can be converted into an immune-activating state. Snyder et al. demonstrate that triggering necroptosis, a form of programmed inflammatory cell death, releases potent Damage-Associated Molecular Patterns (DAMPs) that activate dendritic cells [11]. Chen et al. showed that cellular senescence in cancer cells can trigger a specialized secretome (SASP) that recruits and activates CD8+ T-cells to the tumor site [20]. Zimmermannova et al demonstrated that forced expression of transcription factors PU.1, IRF8 and BATF3 was sufficient to induce the cDC1 phenotype in 36 cancer cell lines of human and mouse origin [13]. Linde et al. showed that myeloid-lineage reprogramming can convert cancer cells into tumor-reprogrammed antigen presenting cells which can then be used for cancer vaccination [12]. Sasso et al. showed that lethally irradiated tumor cells expressing VEGF-C, when injected intradermally as an adjuvant, can stimulate lymphangiogenesis and promote T-cell activation and tumor control in murine melanoma models [10]. Our findings are distinct in many respects from the above studies. Our present study is a follow up to our prior study that shows the live fraction of B16-Ova cells generated after etoposide treatment (but not the dead cell fraction) is immunogenic. Furthermore, in this study, we show that the live etoposide-treated B16-Ova fraction is not beta-galactosidase-positive in the window where T-cell activation is observed in our co-culture assay. Lastly, B16-Ova cells used in our study are not engineered to express transcription factors or growth factors. Elucidating the molecular pathways involved will reveal potentially new mechanisms by which live etoposide-treated B16-Ova cells promote dendritic cell and T-cell activation.

## ACKNOWLEDGEMENTS

We thank Dr. Jennifer Lanni, Dr. Juvenal Lopez, Dr. Robert Morris, Elizabeth Lillie, Erin Post and Heather Anderson for their support and helpful discussion. This project was supported by honors thesis funds awarded to GS, Wheaton Outstanding STEM Scholars (WOSS) Program award to GS and faculty professional development funds awarded to HG and JB.

